# Portable quantum-sensor magnetomyography decodes fine hand movements

**DOI:** 10.64898/2025.12.23.696299

**Authors:** Antonino Greco, Thomas Middelmann, Carsten Mehring, Justus Marquetand, Markus Siegel

## Abstract

Optically pumped magnetometers (OPMs) are compact quantum sensors that can measure the magnetic fields generated by muscle activity (magnetomyography, MMG) without skin contact. This contact-free alternative to surface electromyography (EMG) has remained mostly confined to magnetically shielded rooms and simple tasks, limiting translation. Here, we show that a portable OPM-MMG system operated inside a compact magnetic shield can robustly decode fine finger movements and recover EMG-like muscle activation patterns. Eight participants executed flexion-extension combinations spanning 15 finger actions while we recorded triaxial MMG and bipolar EMG concurrently. MMG supported robust multi-class and finger-specific decoding, recovering a representational geometry that closely matched EMG. Orientation analyses showed that components orthogonal to the muscle axis contributed most significantly to discriminability, providing actionable guidance for OPM array design. Our results show that OPM-MMG in a portable shielded environment preserves task-relevant neuromuscular information and approaches EMG-level structure without skin contact or a magnetically shielded room. Our findings open a path toward hygienic, rapid-setup assessment and human-machine interfacing in clinics and rehabilitation settings.

## Introduction

Noninvasive monitoring of muscle activity is central to clinical diagnostics, neurorehabilitation, and human–machine interfaces. The current standard, surface electromyography (EMG), measures muscle electrical signals via electrodes on the skin. EMG is powerful but requires careful electrode placement, skin preparation, and can be affected by noise at the electrode–skin interface. A complementary approach is to measure the magnetic fields generated by muscle electrical currents – a technique known as magnetomyography (MMG) (*1*–*3*). Modern optically pumped magnetometers (OPMs), which are compact room-temperature quantum sensors, enable sensitive, contact-free detection of these muscular magnetic fields (*4*). Because magnetic fields are less distorted by surrounding tissues and are not affected by impedance variations at the skin, MMG can provide a cleaner or more direct window into muscle activity under certain conditions (*1, 3, 5*–*9*). Indeed, recent advances in OPM technology have allowed researchers to record bio-magnetic signals from the brain, heart, peripheral nerves, and skeletal muscles with high fidelity (*3, 7, 9*–*18*). Triaxial OPM sensors can measure all three orthogonal magnetic field vector components simultaneously, which is advantageous for capturing the full complexity of neuromuscular activity. Such capabilities open new opportunities for gesture recognition and control of assistive devices: for example, contact-free MMG could be exploited in human–machine interfaces (HMIs), prosthetic limb control, and neuromuscular diagnostic tools (*3, 9, 19*–*25*).

Despite this promise, most MMG studies to date have been constrained to highly controlled settings. Many demonstrations have been conducted in large magnetically shielded rooms or have focused on very simple, single-finger movements in individual subjects (*12, 19, 26*–*28*). These requirements limit the real-world applicability of MMG for clinical or assistive technologies. Key questions remain unresolved: Can OPM-based MMG generalize to more complex, fine-grained hand movements? Can it operate reliably in a compact, portable magnetic shielding enclosure that would be practical in a hospital or rehabilitation clinic? And importantly, does the muscle activation information captured by OPM-MMG resemble that of traditional EMG, given the differences in signal type and noise?

Here, we address these questions by combining an OPM array with a portable magnetic shield to record forearm muscle activity during a rich set of finger movements. We asked eight participants to perform 15 distinct combinations of finger flexion and extension while simultaneously recording muscle signals using both OPMs and standard EMG electrodes. All measurements were performed inside a lightweight cylindrical 4-layer shielding tube rather than a full shielded room, bringing the setup closer to a deployable form factor. We quantified the signal quality of OPM-MMG versus EMG, compared the multivariate pattern structure of muscle activity between the two modalities, and evaluated how well each modality could decode (classify) the performed movements. In addition, we analyzed the contributions of each magnetic sensor axis. Our study demonstrates that portable OPM-based MMG can decode complex hand movements and captures muscle activation signatures that closely parallel those of EMG, highlighting its potential for noninvasive neuromotor interfaces.

## Results

We collected data from eight healthy adult participants performing 15 instructed finger movement combinations (all possible combinations of index, middle, ring and little finger flexion-extension). Figure 1 illustrates the experimental setup and task design. Each trial began with a visual cue on a screen indicating a specific combination of fingers to flex (Fig. 1D), and the subject executed that movement at the go cue. All recordings took place inside a compact, cylindrical four-layer mu-metal shield with open ends, allowing the participant’s forearm to be comfortably placed inside (Fig. 1B, C). Crucially, no magnetically shielded room was used; the portable shield provided a local low-field environment for the OPM sensors. We positioned an array of nine triaxial OPMs around the forearm to measure the magnetic signals from the underlying muscles, and we placed nine bipolar EMG electrode pairs on the skin over the forearm muscles (Fig. 1A). This setup enabled the simultaneous recording of muscle activity with both modalities for direct comparison.

**Figure 1:**
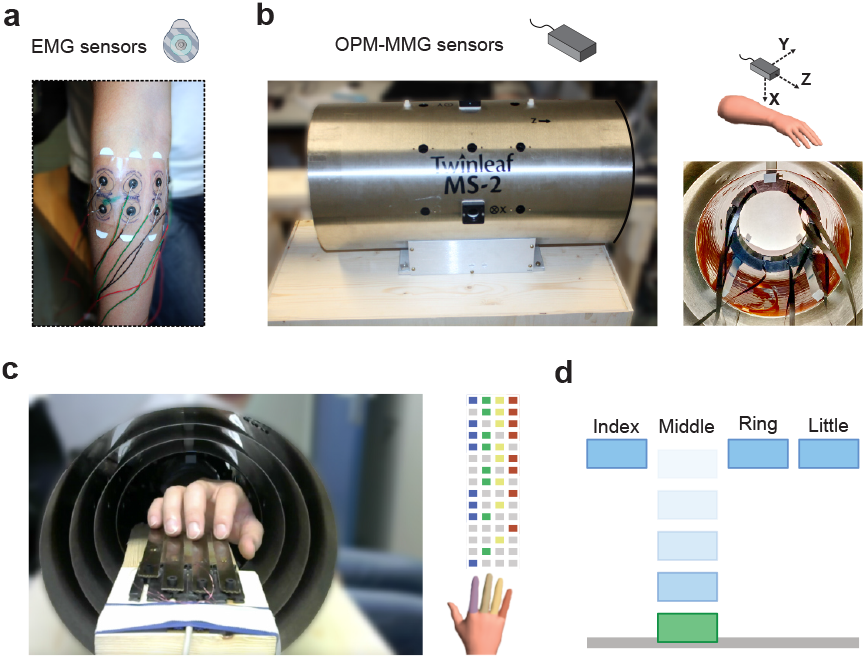
Experimental settings and design. **(a)** Arrangement of bipolar EMG electrodes on an example participant. **(b)** left: lateral outside view of the mobile magnetic shield tube; right bottom: inside view along the axis of the shield tube; right top: illustration of the axis of the three axes of the OPM sensors relative to the forearm. **(c)** left: experimental setup for the finger movement task with 4 buttons recording the finger movement; right: illustration of the 15 combinations of finger movement in the task. **(d)** Illustration of visual cues for the finger movement task. For each combination and trial, the corresponding rectangles moved downwards and finally hit the gray horizontal bar, which signaled the time to perform the corresponding finger movement.

We first examined the raw and averaged muscle responses to confirm that OPM-MMG signals were indeed capturing the finger movement activity. After band-pass filtering (25-100 Hz) and Hilbert envelope extraction, both EMG and OPM-MMG clearly showed phasic increases time-locked to finger flexion onsets. Figure 2A shows grand-average EMG and OPM-MMG signal envelopes for two example movements (flexion of the index finger vs. the little finger) across all participants. Both modalities picked up the activation of forearm muscles, with EMG generally exhibiting a higher amplitude and clearer onset compared to OPM-MMG. The spatial pattern of activation (which sensors/electrodes responded most strongly) was similar between EMG and MMG, as visualized by projecting the average signal amplitudes onto a forearm model (Fig. 2A, bottom). However, the OPM traces appeared noisier than EMG, which is expected, given that magnetically measured signals include more environmental noise.

**Figure 2:**
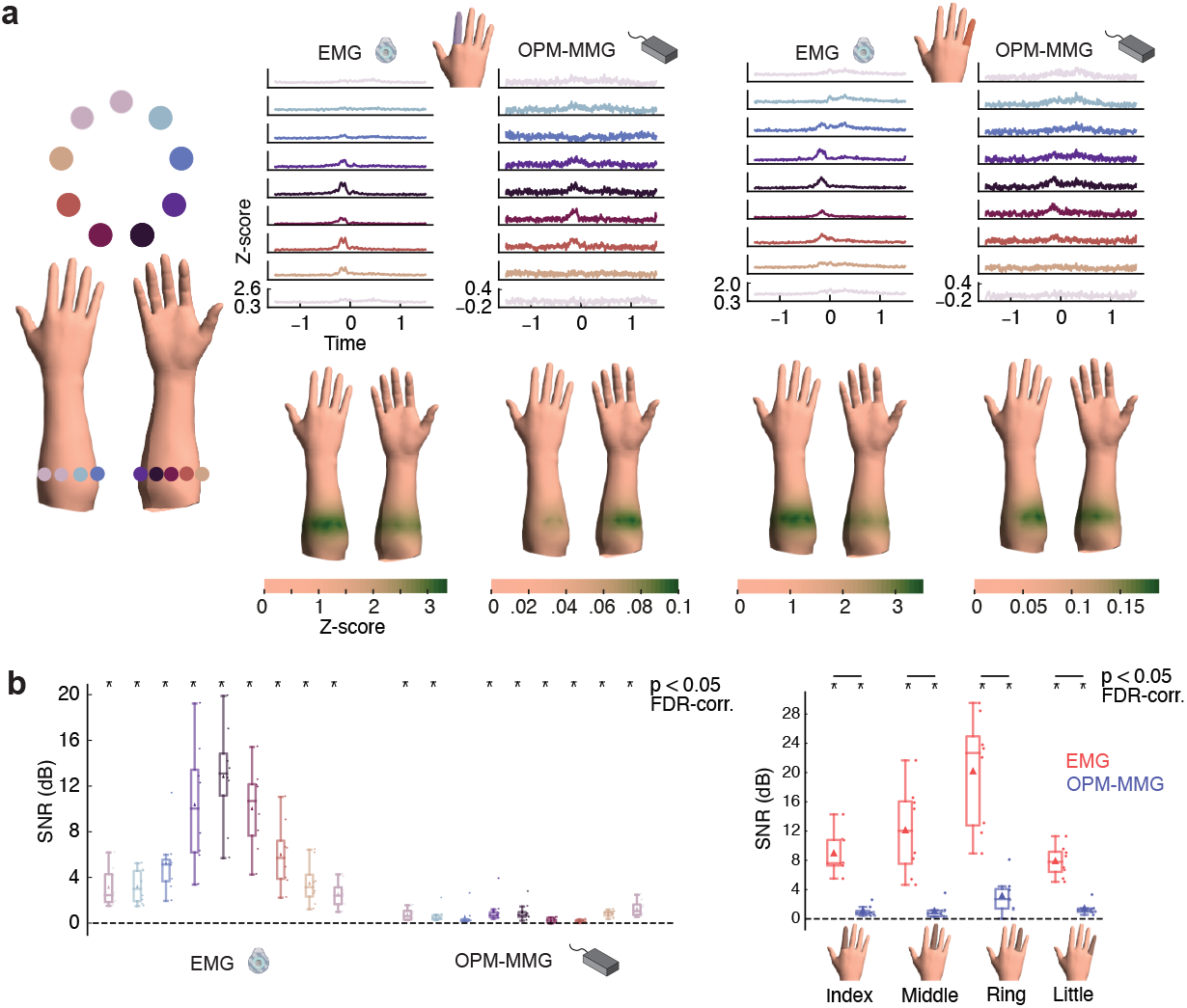
EMG and OPM-MMG signals. **(a)** left: schematic layout of sensor positions on the forearm; right: grand average EMG and OPM-MMG signals across all subjects for index (left) and little (right) finger movements (first principal component for OPM-MMG signals); bottom: average signal between −0.5 and s around movement onset projected on the forearm surface using a template 3D model. **(b)** left: signal-to-noise ratio (SNR) for EMG and OPM-MMG averaged across all movement combinations for each sensor; right: SNR of both sensor modalities averaged across all movement combinations, which required each of the four fingers to be moved, and taking the maximum SNR across sensors.

To quantify signal quality, we computed the signal-to-noise ratio (SNR) for each sensor of each modality. SNR was defined as the ratio of the root-mean-square signal amplitude during the movement (0 to 0.5 s after cue) to that during a baseline period before movement, expressed in decibels (see Materials and Methods). Both modalities yielded significant SNR above 0 dB for muscle activation (Fig. 2B left). Across all 15 movements, EMG sensors achieved a higher average SNR than OPM sensors (p < 0.05 corrected, for each sensor pair comparison). Nonetheless, the majority of OPM sensors also showed clear above-baseline signals (SNR > 0, p < 0.05) for the finger movements. Figure 2B (left) shows the distribution of SNR values for all sensors, highlighting that while EMG had superior SNR overall, the OPM-MMG still reliably detected muscle activity. When considering the best sensor for each individual finger movement (Fig. 2B, right), EMG again had higher SNR (p < 0.002 corrected, for each finger movement), but importantly, OPM-MMG yielded significant positive SNR (p < 0.032 corrected) for every individual finger movement tested, confirming that even fine single-finger actions were discernible in the magnetic recordings.

We next asked whether the pattern of muscle activity across different movements was represented similarly in OPM-MMG and EMG. In other words, do the two modalities “see” the relationships between different finger movements in the same way? To assess this, we performed a pattern similarity analysis. For each modality, we computed a 15×15 dissimilarity matrix reflecting how distinguishable the multivariate muscle activation patterns were between every pair of movements (using cross-validated Mahalanobis distance as the dissimilarity metric; see Methods). Figure 3A (bottom) shows the group-averaged dissimilarity matrices for EMG and OPM-MMG. Visually, the two matrices appear highly alike – for example, certain finger combinations (such as those involving adjacent fingers) were similarly confused or similar in both modalities. We projected these dissimilarity structures into two dimensions with multidimensional scaling (MDS) for visualization (Fig. 3A, top). The EMG and OPM points form very comparable configurations: movements that cluster together in EMG space also cluster together in OPM space, and overall, the geometric arrangement of movement types is preserved between modalities. To quantify this correspondence, we calculated the correlation between the dissimilarity values of EMG and OPM-MMG across all movement pairs, correcting for noise and limited sample size (using Spearman’s attenuation correction). The resulting noise-corrected correlation was extremely high (r = 0.931) and not significantly different from 1 (p = 0.64), indicating that OPM-MMG captured essentially the same representational structure of muscle activity as EMG (Fig. 3C). In practical terms, this means the way muscles activate for various hand movements, and the similarities or differences between those activation patterns, were mirrored in the magnetic signals recorded by OPMs.

**Figure 3:**
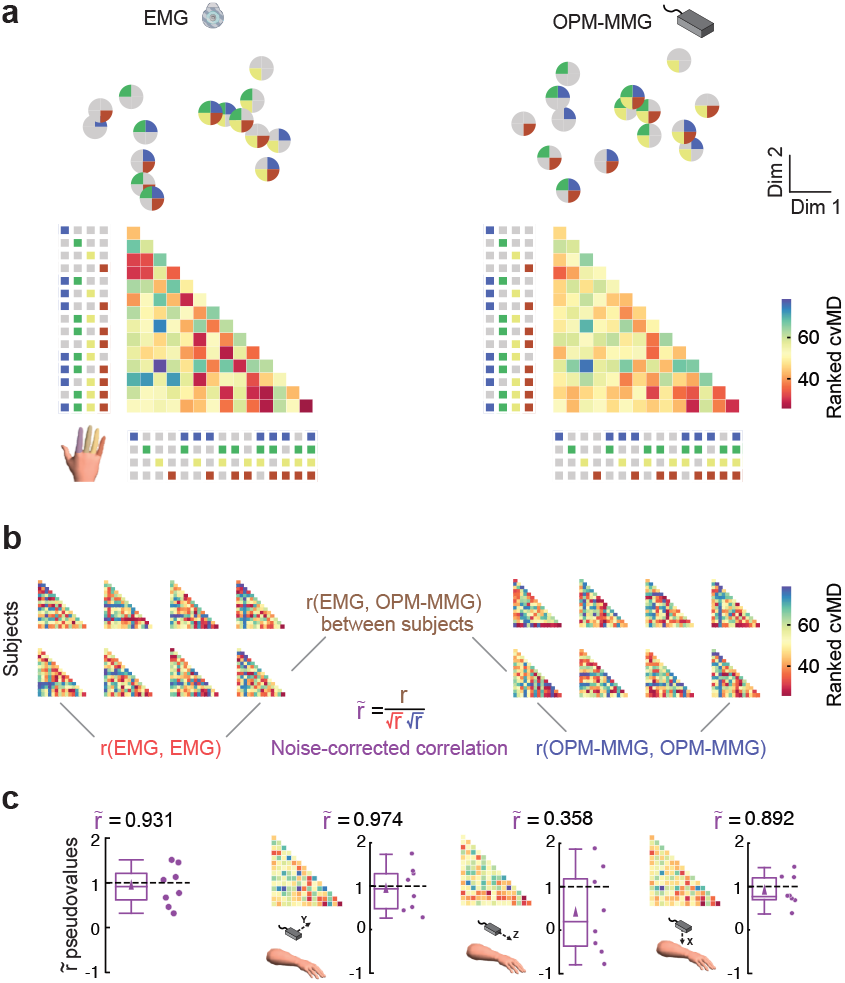
Pattern similarity analysis (PSA) between EMG and OPM-MMG. **(a)** bottom: dissimilarity matrices showing the ranked cvMD values between each combination of finger movements for EMG and OPM-MMG sensors; top: procrustes-aligned Multidimensional Scaling (MDS) projection of the dissimilarity matrices. Each dot represents one movement combination, with each color indicating the movement of a specific finger. **(b)** Illustration of the methodology to derive noise-corrected correlations between EMG and OPM-MMG statistical structures underlying finger movements. The dissimilarity matrices shown for each sensor modality are the single-subject estimates used for group-level comparison. **(c)** Boxplots of the noise-corrected correlations and their respective pseudovalues for the EMG and PCA-reduced OPM-MMG data comparison and for each axis independently. For single axes, the dissimilarity matrix is shown next to the boxplots.

We also examined whether all three OPM sensor axes contributed equally to this similarity structure. Our OPMs measure the magnetic field in orthogonal directions (two roughly orthogonal to the forearm surface, denoted x- and y-axes, and one roughly parallel, z-axis). In the above analysis, we condensed the triaxial OPM data for each sensor into a single combined signal (first principal component across axes) to simplify the comparison. When we instead computed separate dissimilarity matrices for each axis, we found that the two orthogonal axes individually still showed high correspondence with the EMG pattern (r = 0.892, p = 0.408 and 0.974, p = 0.696 for x and y, respectively) and were not significantly different from the EMG structure. The parallel z-axis, however, showed a lower correlation with EMG (r = 0.358) and a trend toward less correspondence (p = 0.125). This suggests that the orthogonal field components measured by the OPMs carry most of the informative signal that aligns with EMG, whereas the parallel component is less informative, possibly due to how the forearm musculature’s magnetic field projects outward.

Finally, we tested how well we can decode which movement was performed using each sensor modality. We trained multiclass classifiers to predict the finger movement combination on each trial from the pattern of either EMG or OPM-MMG activity. We used a nearest-centroid classification approach with cross-validation (five-fold stratified) and evaluated performance as balanced accuracy (chance level = 6.7% for 15 classes; see Methods for details). Figure 4A summarizes the 15-class decoding accuracies. Both modalities allowed above-chance classification of the 15 movements. EMG had significantly higher accuracy than OPM-MMG (p < 0.001), consistent with its higher SNR, but importantly, OPM-MMG’s performance was well above chance (p = 0.007). Moreover, individual participants who were easier to decode with EMG tended to also be easier with OPM: across subjects, EMG and OPM accuracies were strongly correlated (r = 0.78, p = 0.021). At the single-subject level, 6 out of 8 participants showed OPM-MMG decoding accuracy significantly above chance (permutation test, α = 0.05), compared to 7 out of 8 for EMG. These results demonstrate that even without any skin electrodes and in a portable setup, OPM-MMG can discriminate many fine finger movement classes, though with some performance gap relative to EMG.

**Figure 4:**
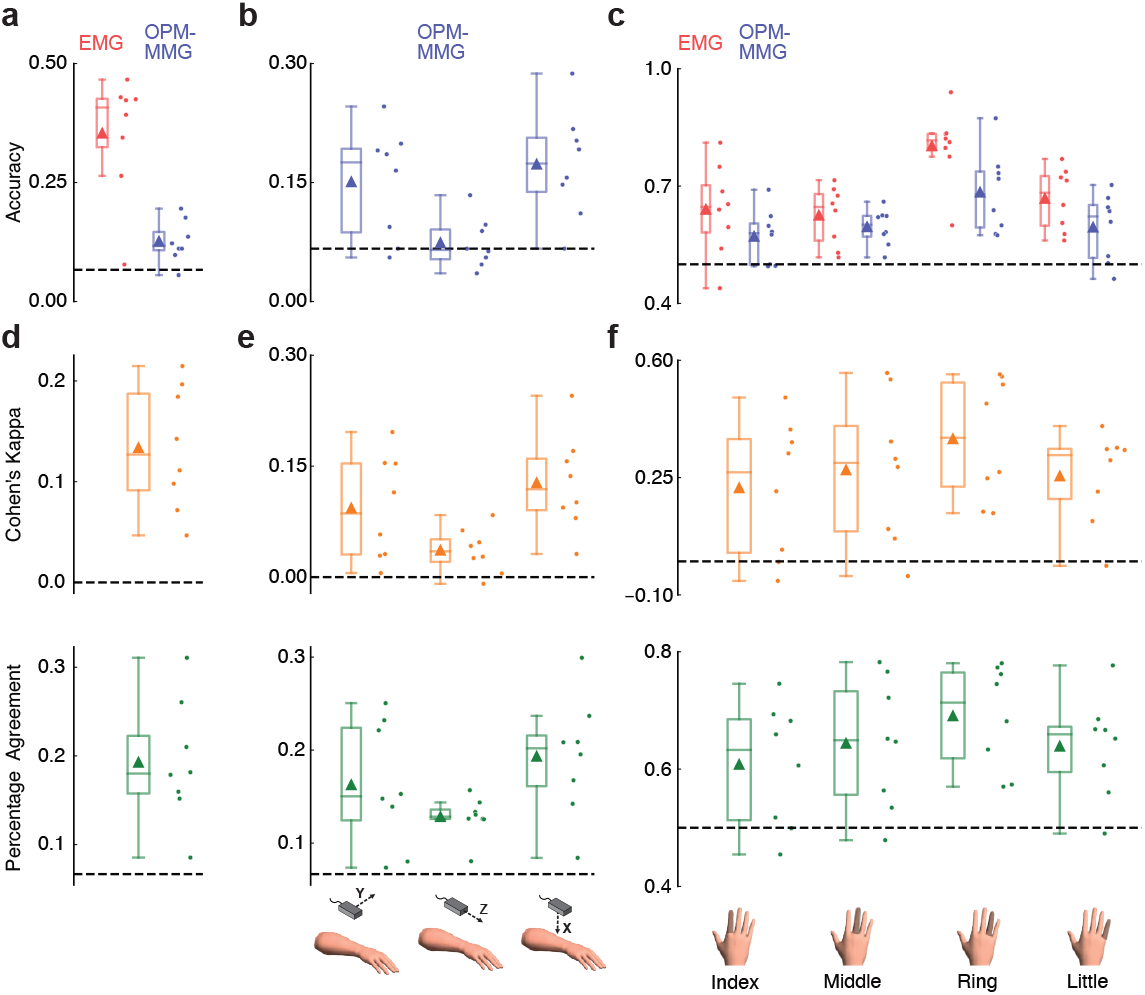
Classification of finger movements from EMG and OPM-MMG. **(a)** Balanced accuracy scores for EMG and OPM-MMG for the classification of all combinations of finger movements. **(b)** Balanced accuracy for each axis of OPM-MMG. **(c)** Balanced accuracy scores for EMG and OPM-MMG for individual finger movement classification. **(d)** Cohen’s kappa and percentage agreement scores between EMG and PCA-reduced OPM-MMG for the classification of all finger combinations. **(e)** Cohen’s kappa and percentage agreement scores for all combination classification between EMG and individual OPM-MMG axes **(f)** Cohen’s kappa and percentage agreement scores between EMG and OPM-MMG for individual finger movement classification. Horizontal dashed lines indicate chance level.

We also examined decoding of individual fingers. Here we reformulated the task as four separate binary classifications: for each finger (index, middle, ring, little), classify whether that finger was involved in the movement or not (regardless of what other fingers moved). This is a relevant scenario for prosthetic or orthotic control, where one may want to detect the intention to move a particular finger. Both OPM-MMG and EMG achieved high accuracy on these single-finger detection tasks (Fig. 4C). Across participants, all four fingers could be detected significantly above chance by both modalities (p < 0.05 for each, FDR-corrected). EMG was slightly more accurate than OPM for certain fingers, significantly so for the little finger (p = 0.048, corrected) and trending for the ring finger (p = 0.066), but for index and middle fingers the difference was not significant. Importantly, the movement predictions made by the OPM-based classifier were consistent with those made by the EMG-based classifier. We quantified this agreement using Cohen’s kappa and percent-agreement metrics: for both, the 15-class classification and each single-finger classification, the OPM and EMG predictions had a mean kappa and agreement above chance (all p < 0.025, corrected) (Fig. 4D–F). This indicates that OPM-MMG and EMG were making similar errors and correct decisions on a trial-by-trial basis, reinforcing that they rely on the same underlying muscle activation information.

Finally, we assessed the contribution of different OPM axes to decoding. We repeated the 15-class classification using only a single axis of the OPM sensors at a time. Consistent with the pattern analysis results, we found that the two orthogonal axes (x and y) of the OPM array were each sufficient to decode the finger movements above chance (p < 0.02, corrected for each), whereas the radial z-axis alone did not perform above chance (p = 0.56, corrected) (Fig. 4B). When using only one axis, performance naturally dropped compared to using all axes together. However, the classifier predictions from the individual OPM axes remained significantly correlated with the EMG predictions (all p < 0.012, corrected).

## Discussion

In this study, we show that portable quantum-sensor magnetomyography can capture robust and informative muscle signals for decoding fine hand movements. Using an array of zero-field OPMs in a compact mobile magnetic shield, we recorded forearm muscle activity sufficient to discriminate 15 distinct finger movement combinations at the group level. Despite lower signal amplitude and higher noise than EMG, the multivariate structure of the OPM signals was virtually indistinguishable from simultaneous EMG, indicating that contact-free MMG preserves the essential neuromuscular information required to tell different movements apart.

Our work extends previous magnetomyography studies (*19*) in three ways. First, we move from single-movement, often single-subject demonstrations to a combinatorial set of finger actions studied at the group level, showing that OPM-MMG supports fine-grained decoding in a behaviorally relevant action space. Second, we translate measurements from magnetically shielded rooms to a portable shielding device, narrowing the gap to clinical and rehabilitation environments (*29*–*31*). Third, by formalizing the representational similarity to EMG and dissecting axis-specific contributions, we provide principle information for designing future MMG sensor arrays.

These results have immediate implications for bioengineering and neural interfaces. Contactless sensing can simplify preparation, avoid skin irritation, and improve hygiene, which is advantageous for repeated use in clinics and rehabilitation. A portable OPM-MMG device could monitor muscle recruitment in patients with motor impairments or provide control signals for prosthetic and assistive devices when EMG is impractical or unstable. Our findings suggest that MMG-based decoding can serve as an alternative or complement to EMG in noninvasive neuromotor interfaces and could be integrated with other modalities to increase robustness.

The orientation analysis yields guidance for next-generation MMG systems: sensors oriented orthogonal to the muscle carry most of the discriminative information, whereas parallel components contribute less. This likely reflects the geometry of muscle current loops and indicates that wearable MMG sleeves should prioritize orthogonal sensitivity and coverage over relevant muscle compartments.

Several limitations should be acknowledged. We studied a modest sample of healthy adults and a constrained, paced finger-press task. Generalization to unconstrained, continuous movements, to different limb segments, and to clinical populations remains to be established. Our setup still relied on a local magnetic shield; operation in fully unshielded environments will likely require active field compensation (*32*). Finally, we used a limited number of sensors in a fixed configuration and tested within-session decoding only, so the optimal sensor layouts and long-term stability of OPM-MMG remain open questions.

Future work should increase sensor density and coverage, integrate active field stabilization, and explore more powerful decoding methods to improve performance. Tests of between-subject generalization, longitudinal stability, and closed-loop control will be crucial to assess translational potential. Overall, our results establish that a portable OPM-based MMG system can achieve EMG-like decoding of fine finger movements, positioning quantum-sensor MMG as a practical tool for neuromuscular assessment and human-machine interfacing beyond traditional shielded facilities.

## Acknowledgements

This research was supported by the European Research Council (AdG 101055186 to O. Röhrle), by the German Research Foundation (DFG; https://www.dfg.de/) project 548605919 (SPP 2311; to J.M.), by the Federal Ministry for Research, Technology and Spaceflight (BMFTR) through the Cluster4Future QSens (Grant ID: 03ZU2110FD; to M.S. and J.M.) and by the German Space Agency (DLR) through funds provided by the Federal Ministry for Economic Affairs and Climate Action (BMWK) (DLR 50BM2534B; to J.M.). We thank Yannic Ober for his support during the experiments.

## Author contributions

A.G.: conceptualization, software, formal analysis, visualization, writing - original draft, writing - review and editing. T.M.: investigation, resources, writing - review and editing. C.M.: conceptualization, supervision, resources, project administration, writing - review and editing. J.M.: conceptualization, supervision, investigation, resources, project administration, writing - review and editing. M.S.: conceptualization, supervision, resources, project administration, funding acquisition, writing - original draft, writing - review and editing.

## Data availability statement

The data that support the findings of this study are available from the authors upon reasonable request.

## Materials and Methods

### Participants

Eight healthy adult volunteers participated in the study (N = 8). All participants provided written informed consent. The study protocol was approved by the Ethics Committee of the University of Tübingen and conducted in accordance with the Declaration of Helsinki.

### Experimental Setup

All experiments were performed in an ordinary, unshielded laboratory room. To reduce ambient magnetic noise, we used a portable magnetically shielding enclosure (MS-2, Twinleaf LLC) consisting of four nested mu-metal cylinders. We removed both endcaps of the cylindrical shield to allow the participant’s forearm to extend through. A similar setup and the penetrating magnetic field inside the magnetic shield is described in more detail in (*30*). Nine triaxial zero-field OPM sensors (QZFM Gen.3, QuSpin Inc.) were mounted in a 3D-printed circular holder at the center of the shield, arranged around the forearm. Participants sat comfortably and rested their forearm inside the shield on a wooden arm support that was mechanically isolated from the shield. We ensured that neither the arm nor the armrest touched the sensors or shield during recording, preventing vibrational or motion artifacts.

### Behavioral Task

Participants performed a visually guided finger movement task involving all combinations of index, middle, ring, and little finger flexion (15 distinct movement classes). On each trial, a graphical cue on a screen instructed the upcoming finger movement. The cue consisted of four vertical bars representing the four fingers; for a given trial, the bars corresponding to the required finger(s) descended toward a horizontal target line. When the bars hit the target line, it signaled the participant to execute the cued finger movement within a 0.5 s response window. Participants pressed corresponding buttons with the instructed finger(s) to register the movement onset.

All 15 finger movement combinations were performed by each participant in pseudorandom order across 150 trials (10 repetitions of each combination). Each flexion movement was a rapid tap (button press) followed by extension to the resting position. Participants were instructed to move all cued fingers simultaneously and to keep uncued fingers as relaxed as possible. They maintained a consistent movement speed and took short breaks as needed to avoid fatigue.

### Data Acquisition

We recorded muscular magnetic fields using nine zero-field optically pumped magnetometers (OPMs) arranged around the forearm. These sensors (QZFM Gen.3, QuSpin Inc.) are triaxial, self-contained quantum magnetometers operating in the spin-exchange relaxation-free (SERF) regime to achieve ultrahigh sensitivity. Each OPM has a specified noise floor below 23 fT/√Hz in the 3–100 Hz frequency band and a nominal bandwidth of 135 Hz. The atomic vapor cell inside the sensor is positioned only 6.2 mm from the sensor’s outer surface, allowing an equally small standoff distance from the skin. For operation, the zero-field OPMs require a near-zero ambient magnetic field; this was ensured by the mu-metal shielding (see Experimental Setup) and by compensation coils inside each sensor that can cancel residual fields up to 50 nT. Each OPM outputs three orthogonal magnetic field readings by using two perpendicular laser beams and modulating the field at 923 Hz along each axis, with phase-sensitive lock-in demodulation (*33*). The output signals are inherently dispersive, introducing a slight nonlinearity (≤5% deviation) within the sensor’s ±1 nT dynamic range.

Surface electromyography (EMG) was recorded concurrently to provide a reference for muscle activity. We placed nine bipolar EMG electrode pairs on the forearm over the major flexor muscle groups (approximately co-localized with the OPM sensor positions. All EMG signals were recorded synchronously with the OPM outputs using a GES400 EEG amplifier (EGI, Inc./Philips-Neuro, Eugene). The EMG and OPM channels were thus recorded in parallel, allowing direct comparisons between modalities. EMG and OPM data were recorded with 1 kHz sampling rate.

### Data Preprocessing

All signal processing was performed using Python 3.7. The raw OPM (9 sensors × 3 axes) and EMG (9 bipolar channels) time-series were first demeaned (zero mean) and then band-pass filtered between 25 Hz and 100 Hz (zero-phase fourth-order Butterworth filter). Power line interference was removed by applying a 49 to 51 Hz notch filter (fourth-order Butterworth). Next, we extracted the signal envelope by applying a Hilbert transform to each channel and taking the absolute value of the analytic signal (*19*). The resulting envelopes were downsampled to 200 Hz for analysis. Finally, we segmented the continuous data into epochs from 1.5 s before to 1.5 s after the movement onset (button press) for each trial.

### Signal-to-Noise Ratio (SNR)

For each finger movement combination, we averaged the preprocessed time-series across all its trials to obtain an average response per sensor. Each OPM’s triaxial data were further reduced to a single component via principal component analysis (PCA), and the average sensor responses were projected onto a 3D forearm model with weights decaying exponentially with distance from the forearm surface.

We quantified signal strength relative to noise by defining SNR as the ratio of root-mean-square (RMS) signal amplitude during movement vs. baseline. Specifically, we computed the RMS of each channel’s envelope in the active window (−0.5 to +0.5 s around movement onset) and in an equal-length baseline window (−1.5 to −0.5 s before onset). The SNR was then calculated as 20*log10(active RMS / baseline RMS), yielding a value in decibels for each sensor and condition.

### Pattern Similarity Analysis

We used a pattern similarity analysis to compare the multivariate activity structures captured by OPM-MMG and EMG. For each participant and each modality, we computed a 15×15 dissimilarity matrix across the 15 finger movement conditions using cross-validated Mahalanobis distance (cvMD) as the metric (*34*). This analysis was implemented with a stratified 5-fold cross-validation scheme to avoid bias. Each dissimilarity matrix was then vectorized (using the lower-triangular entries) to serve as a summary of that participant’s representational structure.

To visualize group-level patterns, we averaged the dissimilarity matrices across participants for each modality and applied multidimensional scaling (MDS) (*35*) to project the group-average dissimilarities into a two-dimensional space. We then used a Procrustes alignment (*36*) to optimally superimpose the OPM and EMG configurations, facilitating direct visual comparison of their pattern geometry.

To quantify the similarity of the EMG and OPM-MMG representations, we correlated their dissimilarity values between modalities. For each participant, we computed a Spearman rank correlation between that participant’s EMG dissimilarity vector and OPM dissimilarity vector, since rank-based correlation reduces sensitivity to scaling differences. However, measurement noise can attenuate these raw correlations, so we applied Spearman’s correction for attenuation (*37, 38*) to better estimate the true underlying correspondence between modalities.

In practice, we first assessed within-modality reliability by calculating the Spearman correlation of dissimilarity vectors across participants within each modality. We then calculated the cross-modal Spearman correlation between EMG and OPM dissimilarities across participants (excluding each participant’s self-comparison). Finally, we obtained a noise-corrected EMG–OPM correlation by dividing the observed cross-modal correlation by the square root of the product of the two within-modality correlations (*37, 38*).

Because this noise-corrected correlation is inherently a group-level measure, we used a jackknife procedure to derive an estimate for each individual (*39, 40*). Specifically, we recomputed the noise-corrected correlation n times, each time leaving out one of the n participants, and used these leave-one-out results to calculate a pseudovalue for each participant. These pseudovalues were treated as independent observations of the EMG–OPM correlation for statistical analysis.

### Classification Analysis

We next evaluated the ability of each modality to decode finger movements via pattern classification. We used a nearest centroid classifier (*41*) with Euclidean distance, implemented separately for the EMG data and the OPM-MMG data. Each trial’s data from −0.5 s to +0.5 s around movement onset were converted into a feature vector by concatenating the time series from all sensors. Classifier models were trained and tested using stratified 5-fold cross-validation.

We performed two classification tasks: a multi-class classification of all 15 finger movement combinations, and a set of binary classifications for individual fingers. In the 15-class scenario (chance accuracy = 6.7%), the classifier’s task was to identify which specific combination of fingers moved on each trial. In the binary finger-specific scenario (four separate analyses, chance = 50%), the task was to detect whether a given finger was involved in the movement or not (e.g., index-finger movement vs. no index involvement). For the multi-class models, we used a prototype-based approach: during training, we averaged the feature vectors of all training trials for each class to form a centroid pattern, and we assigned each test trial to the class with the closest centroid (smallest Euclidean distance). Model performance was quantified using the balanced accuracy (the average of per-class accuracies) to account for the unequal representation of classes.

For OPM-MMG data, we evaluated two feature representations: one using the PCA-reduced sensor signals (one principal component per OPM sensor) and one using all individual OPM sensor axes as separate features. Finally, to compare the consistency of classification outcomes between modalities, we examined the agreement between the EMG-based and OPM-based classifiers. For each trial, we determined whether the EMG and OPM classifiers yielded the same class prediction. From this, we computed the percentage agreement (proportion of trials with identical predictions) and Cohen’s kappa, which measures inter-model agreement beyond chance levels.

### Statistical Analysis

All statistical analyses were two-tailed with a significance level of α = 0.05. We used paired t-tests to compare metrics between the two sensor modalities (e.g., EMG vs. OPM SNR or classification accuracy) and one-sample t-tests to determine if metrics differed from a null expectation (e.g., SNR vs. 0, accuracy vs. chance). Where multiple comparisons were made, p-values were adjusted using the false discovery rate (FDR) method (*42*) to control for Type I errors.

For the noise-corrected correlation analysis of representational similarity, which yields a single group-level correlation, we obtained an approximate value for each participant using a jackknife pseudovalue approach (*39, 40*) as described above. These participant-specific pseudovalues were then used for statistical testing (for example, to compute group means and confidence intervals).

For the classification results, we also assessed significance at the single-subject level with permutation tests. For each participant, we generated 5,000 surrogate datasets by randomly shuffling the trial labels (finger movement classes) and reran the entire classification analysis for each shuffled dataset. This procedure yielded a null distribution of balanced accuracy scores for that participant under the assumption of no true class-related signal. The empirical p-value was defined as the fraction of permutations in which the shuffled-data accuracy met or exceeded the actual accuracy. This nonparametric test provides a robust assessment of whether each participant’s decoding performance exceeded chance levels.

